# SIMIT-seq: a bead-free, scalable microfluidic platform for single cell mRNA sequencing with ultrahigh cell-indexing rate

**DOI:** 10.1101/2025.04.23.650129

**Authors:** Mohan Duan, Wenbing Gao, Guojian Li, Yao Cai, Zequan Zhao, Xuedong Lan, Dayin Wang, Xiaoxing Xing, Yuan Luo

**Affiliations:** College of Information Science and Technology, Beijing University of Chemical Technology, No. 15 North 3rd Ring Rd., Beijing, 100029, China; State Key Laboratory of Transducer Technology, Shanghai Institute of Microsystem and Information Technology, Chinese Academy of Sciences, Shanghai, 200050, China; Center of Materials Science and Optoelectronics Engineering, University of Chinese Academy of Sciences, Beijing, 100049, China

**Keywords:** Single-cell mRNA sequencing, Microfluidic barcoding, Micro-well array, Single-cell indexing

## Abstract

SIMIT-seq is a bead-free, scalable microfluidic platform designed for high-efficiency single-cell mRNA sequencing. Conventional microfluidic-based single-cell RNA sequencing platforms rely heavily on barcoded beads and intricate co-encapsulation schemes, often constrained by double Poisson limitations and the complexities of bead synthesis. In contrast, SIMIT-seq eliminates the need for beads entirely by employing a deterministic, orthogonal barcoding strategy within a two-dimensional micro-well array. This platform achieves an impressive single-cell indexing rate of 96.6% without the need for complex microfluidic operations. Here, we describe the design and fabrication of the SIMIT-seq platform, outline its workflow for transcript capture and library preparation, and demonstrate its application in profiling K562 cells. Our results validate both the high fidelity of molecular barcode immobilization and the system’s capability to support downstream single-cell mRNA sequencing. SIMIT-seq offers a cost-effective and scalable alternative for single-cell transcriptomics, providing a promising foundation for future single-cell omics applications.

## Introduction

Ever since its advent, single-cell transcriptomics analysis has revolutionized the field of biomedical research. By offering resolution at individual cell level for mRNA sequencing for biological/clinical relevant samples, single-cell transcriptomics has been instrumental in unveiling tremendous amount of biological insights across all sub-disciplines in biomedical research^1-5^. The very first work on single cell mRNA sequencing published in 2009 performed such analysis by individually hand-picking target cell one at a time, which is obviously tedious and yield extremely low throughput^6^. Modern single-cell transcriptomics platforms have involved past those early stages and reaching high level of processing throughput and platform integration and automation. There are three major technological strategies for implementing single-cell transcriptomics. The first strategy makes use of flow cytometry instrument for sorting individual cells into well plates and subsequently perform single cell indexing and library construction on those plates^7, 8^. This strategy offers exquisite sensitivity in detecting transcripts from single cell yet only yield relatively low throughput at ~100s cells for each well-plate operation^7^. It is worth noting that robotic pipette manipulation and further plate-level pooling can be used to upgrade the automation and throughput for this strategy^8^. The second strategy relies on the idea of single-cell combinatorial indexing (SCI)^3^ or similarly the split –pool barcoding method^4^, which can reach >100,000 throughput level. However, these techniques still heavily involve tedious manual operation using well plates and have not seen wide adoption in the single-cell research community. The third strategy utilizes microfluidic devices to realize co-encapsulation of single cell and single barcoding beads into micron size compartments, most notably micro-droplets^9, 10^ and micro-well arrays^5, 11^. Cells are then to be lysed within the compartments and mRNAs be released and bound to barcoding DNA molecules on the beads. Hence, every mRNA will be appended with cell-specific barcodes that can be used to index subsequent sequencing reads. The operation of microfluidic devices can be achieved with an integrated and automated apparatus, which makes them primed for wider scale adoption. And it is precisely the microfluidic-based approach that stimulated the implosive development of single-cell transcriptomics by making these intricate technological platforms available to the broader biomedical research community, starting from early vendors like Fluidigm^12^, to the current industry giant 10X Genomics^13^. Notably there is recent effort to increase the cell processing throughput by combining droplet microfluidics with combinatory indexing^14^. More interestingly, microfluidic-free co-encapsulation of cells and beads within micro-droplets has recently been demonstrated using the template emulsification method^15^.

The involvement of barcoding beads is universal for these single-cell microfluidic platforms, albeit certain significant difficulties for the implementation. First, the manufacturing of such single-cell specific barcoding beads relies on the well-known split-and-pool strategy^9^, which requires cumbersome steps of remixing of bead samples during the direct synthesis of DNA molecule on the bead surface. A lesser discussed issue for barcoding beads manufacturing is that, in order to capture mRNA with their polyA tails appended at the 3’ ends, the arrangement of DNA barcode on bead surface needs to be in 5’->3’ direction. This requires non-conventional reverse-direction phosphoramidite synthesis of DNA on bead surfaces, a process that is significantly more costly than typical DNA molecule synthesis (3’->5’ by default)^16^. More critically, the requirement of precise 1:1 ratio co-encapsulation of two micron-sized particles (cells and beads) raised the difficulty of the entire design and implementation of the microfluidic platforms in use. For droplets or micro-well based devices, they are known to face the double Poisson distribution limitation that could drastically decrease the number of cells being precisely 1:1 co-encapsulated with barcoding beads, and therefore fail to be properly indexed^17-19^. A great amount of effort has been spent on overcome such double Poisson limit, including methods like close-packed loading of hydrogel barcoding beads for the droplet-based platforms^20^, DEP-assisted trapping (dTNT-seq^21^) or size-exclusion micro-well design (Well-Paired-seq^17^) for well-based platforms, and hybrid methodologies that involve hydrodynamic trapping followed by active co-encapsulation with droplet-in-oil methods (hydro-seq^19^ and paired-seq^18^). All these ingeniously designed microfluidic devices, albeit reaching fairly high co-encapsulation rate (some higher than 90%), unfortunately complicate the fabrication and operation of the devices significantly.

In this work, we attempt to challenge the basic premise of this strategy and the necessity of the use of barcoding beads. We design a simple micro-well based microfluidic platform (Figure 1), termed Single-cell Indexing via MIcrofluidic-barcoding Technology for sequencing (SIMIT-seq), that is capable of performing highly scalable single-cell indexing without the need for barcoding beads and achieve single cell mRNA sequencing. Most notably, through our method, an impressive 96.6% single-cell indexing rate can be achieved without any complicated microfluidic operations. In the following, we mainly demonstrate the working principle of SIMIT-seq, along with its single cell indexing results, as well as single-cell mRNA sequencing data obtained using the presented platform.

**Figure 1.**
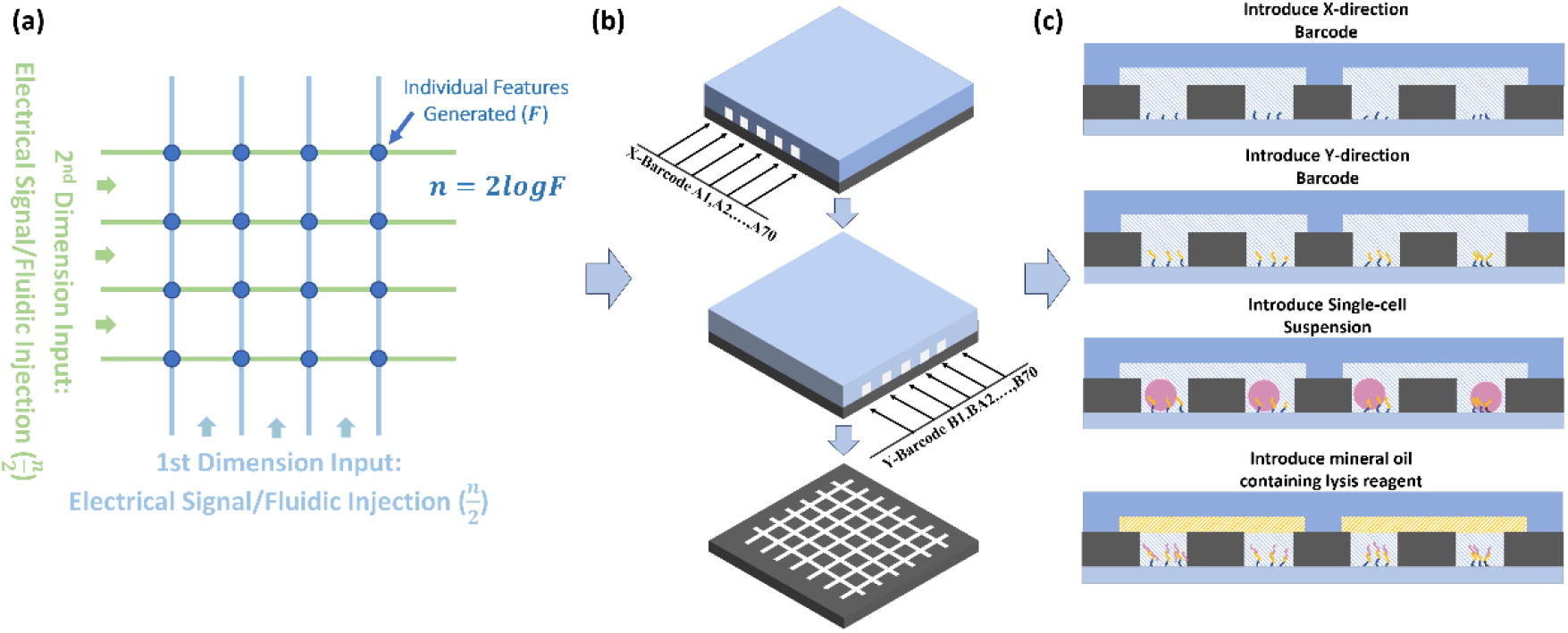
Schematic illustration for SIMIT-seq: (a) the design principle of microfluidic barcoding, (b) creation of 2-dimensional grid using microfluidic channels, (c) experimental procedure for implementing single cell mRNA capturing.

## Materials and Methods

### Device fabrication process

The microfluidic platform comprises a micro-well plate, a pair of orthogonally aligned microchannel devices enabling two-dimensional barcode immobilization, as well as a microchamber for cell loading. The micro-well plate were fabricated on glass substrates using standard photolithography with SU-8 2015 photoresist. The resulted micro-well plate had 4900 micro-wells arranged as a 70 × 70 array, where each well had a diameter of 28 μm and a depth of 25 μm.

The paired microchannel devices for barcoding (channels A and B), as well as the microchamber for cell loading were both fabricated in polydimethylsiloxane (PDMS, Sylgard 184, Dow Corning) using soft lithography. The master molds for PDMS replica molding were all prepared by standard lithography with SU8-2015 on silicon wafers, with photoresist thickness of 30 μm and 60 μm for the barcoding channel and cell loading channel, respectively. PDMS premixed with 10:1 ratio between the gel and curing agent were then poured onto the mold and cured at 60°C for 2.5 h. After curing, these PDMS devices were peeled off from the mold and punched with inlet/outlet holes. Before experiments, the PDMS channel devices were carefully cleaned by isopropanol (IPA) in ultrasonic bath, and were aligned to the micro-well array under microscope to form reversible bonding.

Besides, a PDMS chamber for vacuum-assisted loading of the barcode sample was prepared by replica molding with a laser-cut acrylic block (27.5 mm × 31 mm, and 6 mm thick). An access hole was punched through the chamber ceil for venting the chamber. The complete fabrication workflow is illustrated in Figure 2a.

**Figure 2.**
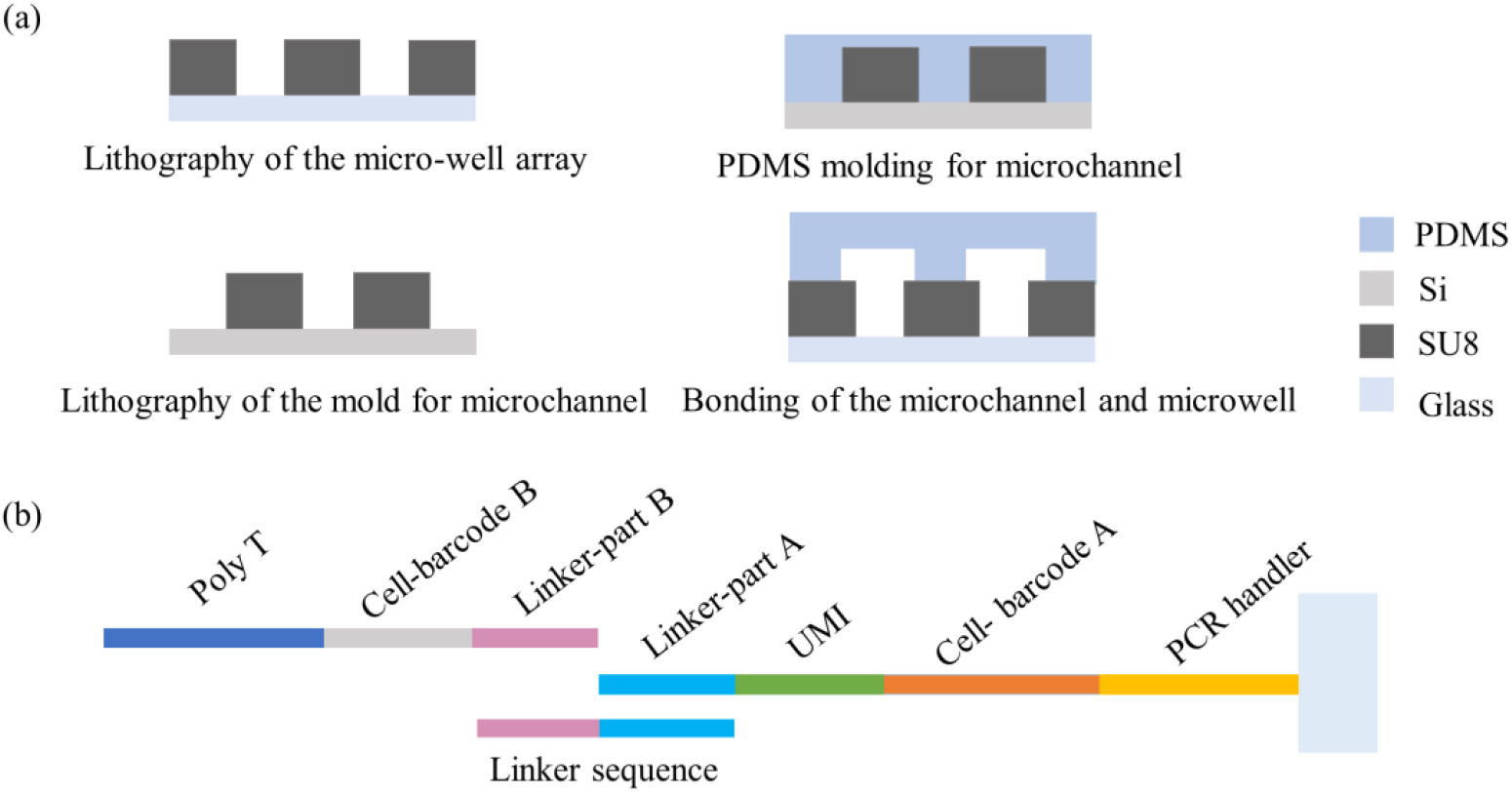
(a) Fabrication process flow for the micro-well plate and the barcoding microchannel. (b) Sequence structure for the barcoding DNA molecules.

### Surface functionalization of micro-well array

To enable covalent attachment of DNA barcodes, aldehyde functional groups were introduced onto the glass surface at the bottom of the micro-wells. The micro-well plate was first cleaned in 1 M KOH by ultrasonic treatment for 15 min, rinsed thoroughly with deionized (DI) water, and dried under a nitrogen stream, followed by baking at 120°C for 10 min. Surface activation was then performed using oxygen plasma (35 W, 15 min), generating hydroxyl groups on the exposed glass. The substrate was immediately immersed in a 2% (v/v) solution of 3-aminopropyltriethoxysilane (APTES) in anhydrous ethanol and incubated in the dark for 2 h. After treatment, the excess silane was removed by ultrasonic washing in ethanol for 5–8 min, followed by multiple DI water rinses and drying under nitrogen. The substrate was then baked at 100°C for 1 h. Thereafter, the aldehyde groups were introduced by incubating the micro-well plate overnight in a 5% (v/v) glutaraldehyde (GA) solution in PBS. Vacuum was applied to the GA container for 10 min at the beginning of the treatment to remove any trapped air in the micro-well plate. After incubation, the micro-well plate was rinsed thoroughly by DI water and stored at 4°C in the dark until further use.

### Barcodes A (x-direction) immobilization

DNA barcode A (from A1-A70), each has a PCR handler, an unique part A of the cell barcode, an unique molecular identifier (UMI), as well as the part A of the linker sequence, were synthesized with a 5’-amine modification (sequences in Table 1). Microchannel A with seventy parallel channels (each was 70μm-width and 30μm apart) was aligned onto the micro-well plate, with each channel overlaid a row of wells in x direction (Figure 1).

**Table 1.**
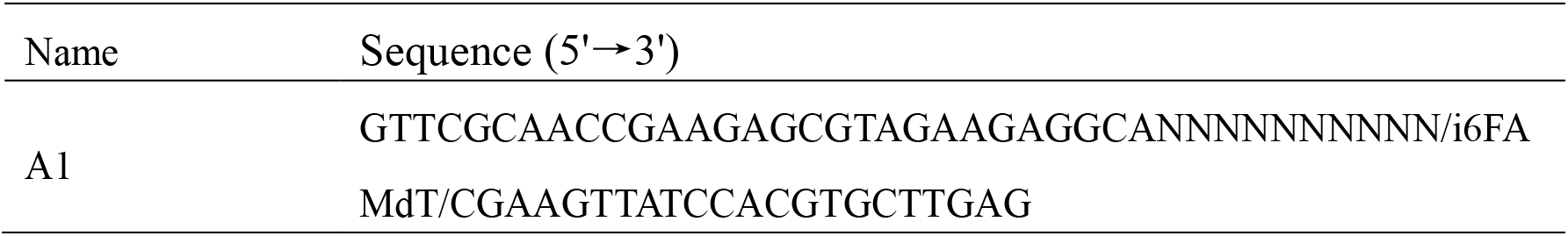
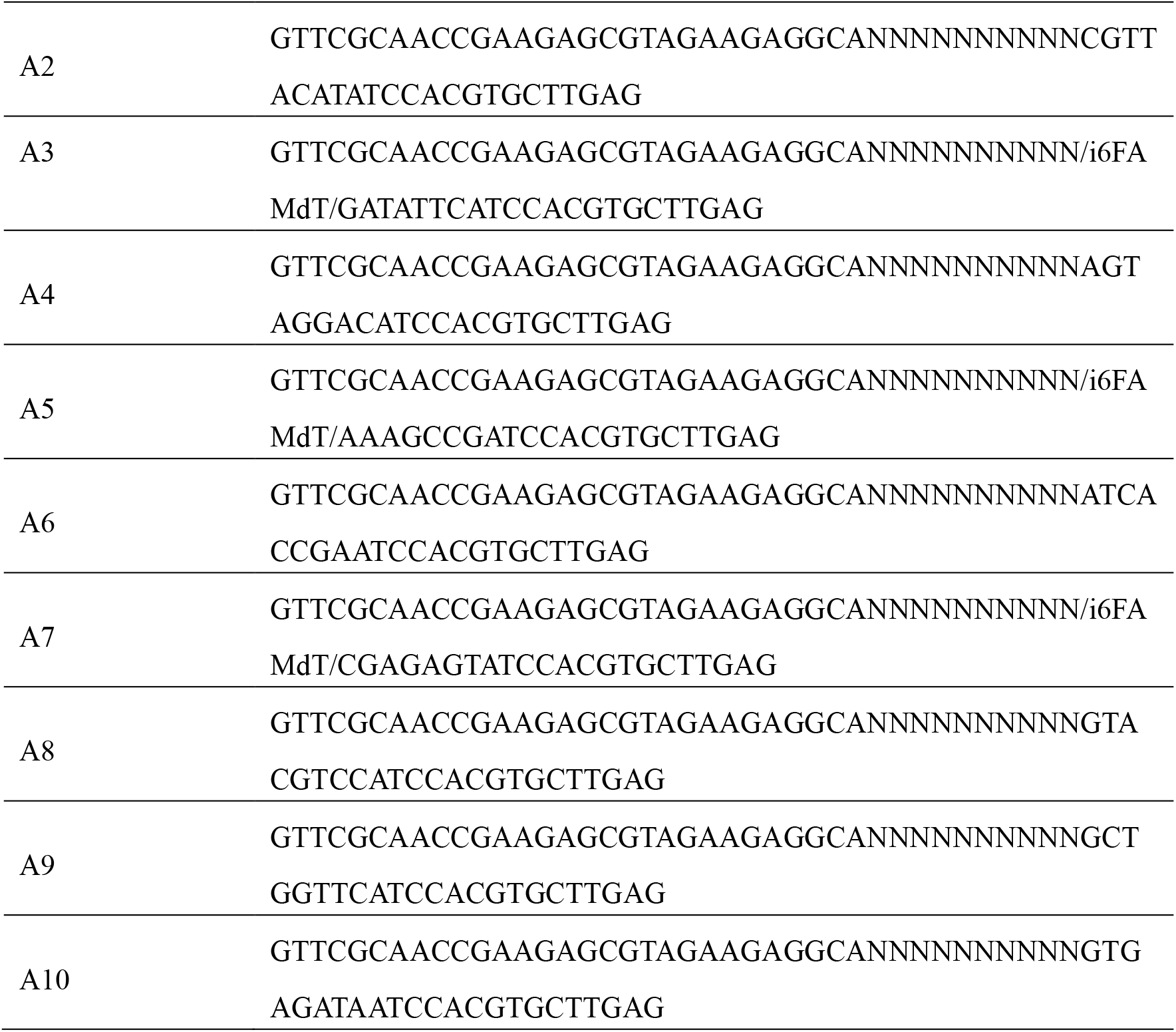
A1-A10 Sequence Structure.

**Table 2.**
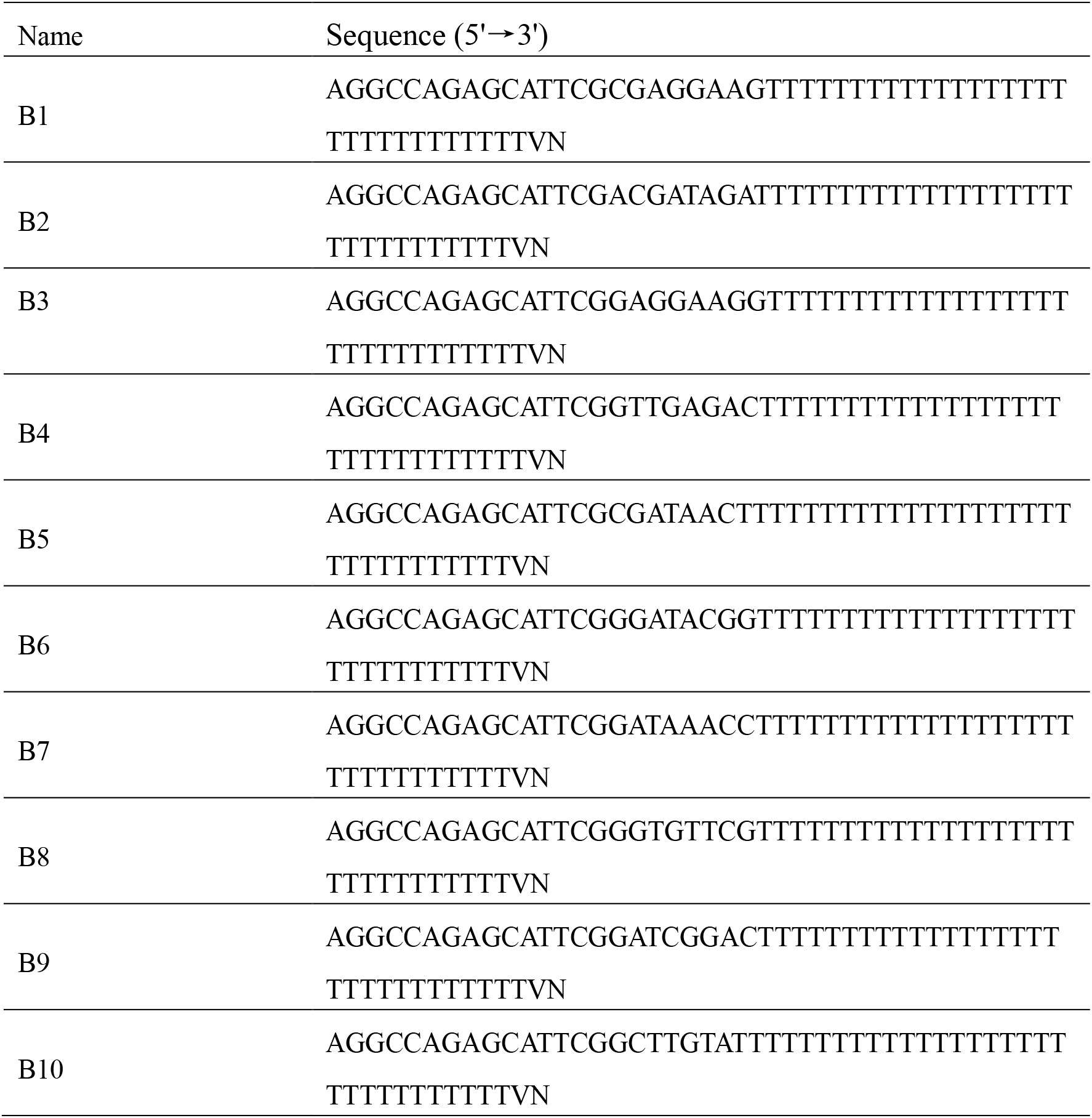
B1-B10 Sequence Structure.

Each barcode A of 25 μM was loaded into the corresponding inlet (5 μL per inlet) of the parallel channels of microchannel A. The PDMS vacuum chamber was fitted over the outlets of the channels, and a negative pressure was applied using a syringe to draw the barcode solutions into the channels and fill the wells. The assembly was incubated in dark on a shaker for 3 h, with periodic vacuum applied to refresh the barcode solution in the channel (every 20 min). The unbound barcode was removed by flushing the channel with 2× SSC buffer after barcode A incubation. The microchannel A was then removed, and the micro-well plate was immediately immersed in sodium borohydride solution (100 mg NaBH_4_ in 37.5 mL PBS, 12.5 mL ethanol, 75 μL Triton X-100) for 15 min to reduce the unreacted aldehydes, and stabilize the Schiff base linkages via reductive amination. The micro-well plate with barcodes A fixed at the bottom of the wells was then rinsed with DI water, dried, and stored in the dark.

### Barcodes B (y-direction) linkage

DNA barcodes B (B1–B70), bearing a 5’-phosphate modification, part B of the linker, an unique part B of the cell barcode, and a poly(dT) tail, were annealed with a linker sequence (Figure 2b) using a slow cooling protocol (95°C for 2 min, then cooled to 20°C at 0.1°C/s). As a result, the part B of the linker of barcode B hybrid with half of the linker sequence, leaving the other half with complementary sequence to the linker (Part A) in the barcode A. A ligation mixture containing 25 μM of the annealed products was then prepared using T4 DNA ligase (400 U/mL) in NEBuffer 3.1 and T4 ligase buffer. Microchannel B featuring 70 parallel channels was aligned orthogonally to the previous microchannel A, such that each channel overlaid a column of wells in y direction (Figure 1b). The ligation mixture (5 μL per inlet) was loaded and drawn into the channels using vacuum-assisted approach as described above. The device was incubated on a shaker in the dark for 3 h, with refresh of the solutions in channels every 20 min. During the incubation, the overhanging part of the linker sequence in the ligation mixture hybridized with the part A of the linker in barcode A, which promoted the alignment of the 5’ ends of barcodes B to the 3’ ends of barcodes A. The 5’ ends of barcodes B then ligate with the 3’ ends of barcodes A under the action of T4 DNA ligase. A schematic diagram of this process is shown in Figure 2b. Following the ligation, the channels were flushed with 2× SSC before being removed. The micro-well plate was then washed with 0.2% SDS in 2× SSC, rinsed with water, and dried with N_2_. A PDMS reservoir was bonded to incorporate the micro-well array to permit subsequent reagent exchange. The linker sequence hybridized to the ligated barcode was removed with 80 mM KOH treatment for 5 min under vacuum. The micro-well plate were rinsed and dried. At this stage, each micro-well contained a unique cell barcode obtained through the ligation of barcode A and B (Ai-Bj), forming a deterministic 2D barcode matrix (Figure 1b).

### Single-cell capture and lysis

Human chronic myelogenous leukemia cells (K562 cells) were exploited for the primary characterization of the developed platform. First, the cell loading chambers with 10 parallel channels was aligned and attached to the micro-well array under microscope, with each channel covering 7 rows of microwells. A custom PMMA clamp was used to strengthen the reversible bonding and thus prevented the leakage issues. The clamped chip was first primed with cell loading buffer, and was then placed in a vacuum chamber for 10 min to remove any trapped bubble. The fetal bovine serum (FBS) solution was subsequently injected into the chip, incubated for 20 min to minimize the cell adhesion to the microchannel walls. After FBS coating, the chip was flushed with PBS solution. Next, the K562 cells co-stained with calcein-AM and Hoechst 33342 were slowly introduced into the chip inlet. The cell flow was carefully controlled by adjusting the height of the tubing connected to the chip outlet. The cells then fall into the micro-wells under gravity in 5 min, and then the remaining cells were gently flushed with PBS. After cell capture, the pre-lysis buffer was slowly injected into the chip. The pre-lysis buffer consists of 160 mM Tris (pH=7.5), 0.5 U/μL RNase inhibitor, and 16 mM EDTA dissolved in 1× PBS.

Meanwhile, mineral oil containing suspended microparticles of surfactant (N-Lauroylsarcosine sodium salt) for cell lysis at 5% (w/v) was prepared, where 0.25g solid surfactant was added into 5ml mineral oil followed by ultrasonic-assisted dispersion for 30 min. For cell lysis, 50 μL of pure mineral oil was first infused into the cell loading chamber to seal the microwells, followed by the introduction of the surfactant-containing buffer. The microparticles of the surfactant sedimented down to the microwells and dissolved in the pre-lysis buffer, leading to rapid cell lysis in the sealed microwells. After cell lysis, the device was incubated at room temperature for 15 min, allowing the PolyA tails of the released mRNA to be captured by the PolyT tail of the barcodes fixed at the well bottom.

### Reverse transcription

After lysis, the micro-well array was washed with 6× SSC (DEPC-treated) and dried. A new PDMS reservoir was aligned and bonded to accommodate the reagents in the following steps. The reverse transcription (RT) mix (Table 3) included Maxima H Minus Reverse Transcriptase, RNase inhibitor, dNTPs, and a template-switching oligonucleotide (TSO; Table 4) to extend transcripts with a defined sequence at the 5′ end. RT buffer was first introduced to prime the wells, followed by the full RT mix. The device was incubated at room temperature for 30 min with gentle shaking, then transferred to a 42°C incubator for 90 min to complete cDNA synthesis.

**Table 3:**
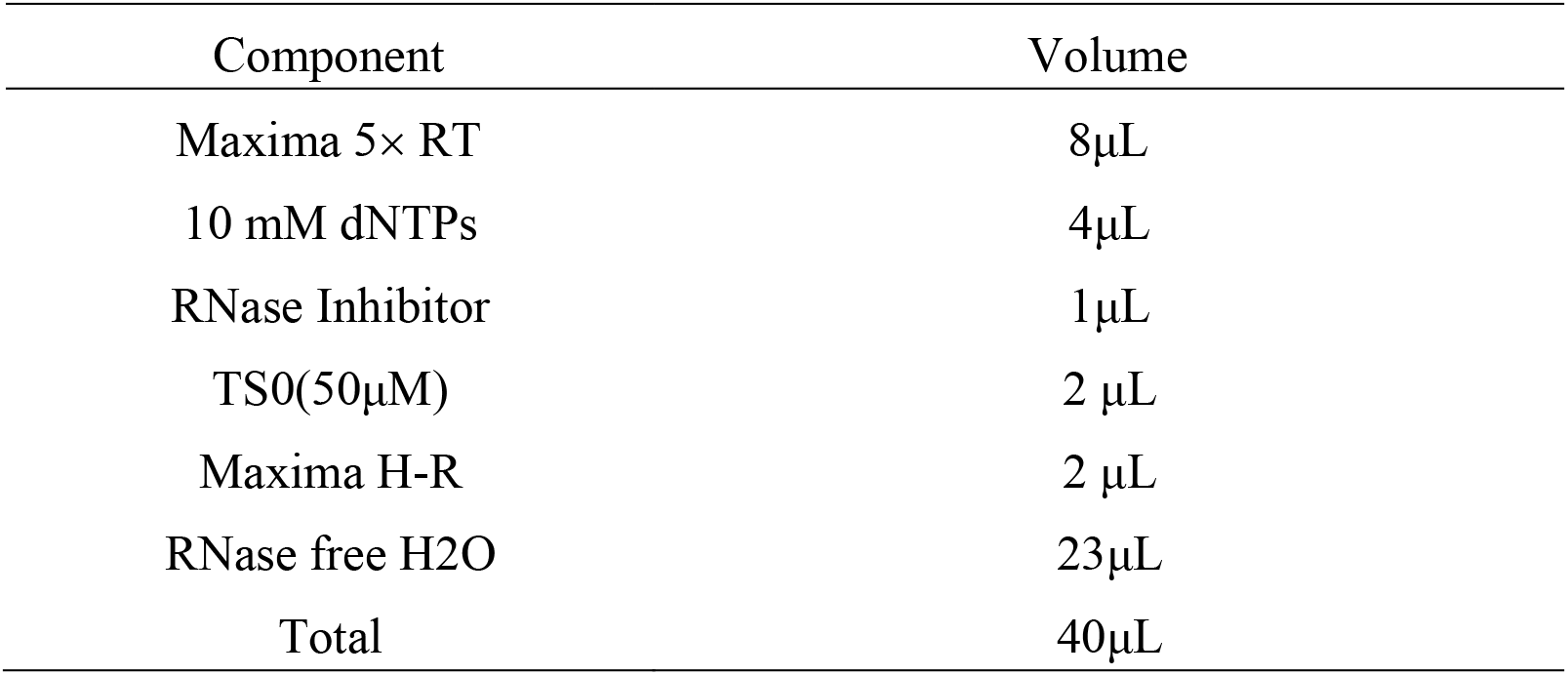
Reverse Transcription Mixture.

| Component | Volume |
| --- | --- |
| Maxima 5× RT | 8μL |
| 10 mM dNTPs | 4μL |
| RNase Inhibitor | 1μL |
| TSO(50μM) | 2 μL |
| Maxima H-R | 2 μL |
| RNase free H<sub>2</sub>O | 23μL |
| Total | 40μL |

**Table 4:**
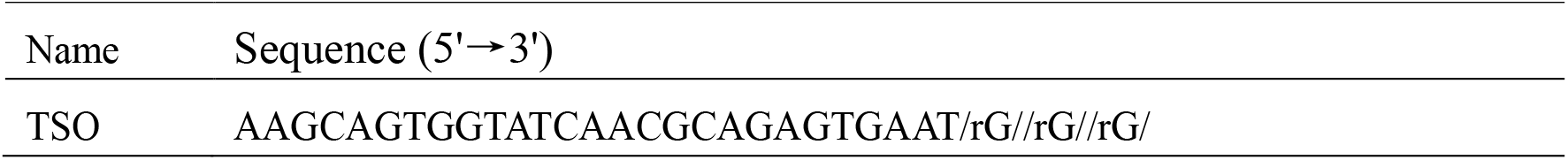
TSO Sequence Structure.

### Second strand synthesis

Following RT, the wells were washed sequentially in TE/TW, TE-SDS, and Tris buffers. Denaturation of cDNA-mRNA hybrids was performed using 80 mM KOH for 5 min to release cDNA into solution. After rinsing and drying, the second-strand extension mix (Table 5), containing NEB Klenow fragment and extension primers, was introduced. The reaction was carried out at 37°C for 2 h.

**Table 5:** Second-Strand Synthesis System.

| Component | Volume |
| --- | --- |
| 10×NEB buffer 2 | 6μL |
| Extension primer ((100μM) | 6μL |
| 10mM NEB dNTP mix | 6μL |
| NEB Klenow fragment | 6μL |
| DEPC-treated water | 36μL |
| Total | 60μL |

### cDNA extraction and PCR amplification

Second-strand cDNA was extracted by incubating the micro-wells with KOH, neutralized with Tris-HCl (pH 7.0), and collected into microtubes. The resulting product, containing full-length barcoded cDNA, was used as template for PCR (Table 6). Reactions were cycled as shown in Table 7: 30 cycles of denaturation (98°C, 20 s), annealing (62°C, 15 s), and extension (72°C, 15 s), following an initial 3-min denaturation at 95°C. After PCR is complete, the samples can be stored at −20°C for later use.

**Table 6:** PCR Amplification System.

| Materials | Volume |
| --- | --- |
| Second strand | 5 µL |
| 10 µM Forward primer | 1.5µL |
| 10 µM Reverse primer | 1.5 µL |
| KAPA HiFi HotStart ReadyMix | 12.5 µL |
| Nuclease-free water | 4.5 µL |
| Total | 25 µL |

**Table 7:** PCR Amplification Program.

| PCR Program |  |  |  |
| --- | --- | --- | --- |
| Pre-denaturation |  | 95°C | 3min |
| 30cycles | Denaturation | 98°C | 20S |
|  | Annealing | 62°C | 15S |
|  | Extension | 72°C | 15S |

### cDNA purification

Amplified cDNA was purified using KAPA Pure Beads with a bead-to-sample ratio of 0.8×. Magnetic separation and ethanol washing were performed, followed by elution in RNase-free water. Purified cDNA was stored at 2–8°C (short term) or −20°C (long term) prior to library preparation. The purified cDNA can be stored at 2°C to 8°C for 1-2 weeks or at −15°C to −25°C for 1 month.

### Tn5 library construction and sequencing

A transposase-based library preparation was performed using the TruePrep® DNA Library Prep Kit V2 (Vazyme). Custom primers P5 and N701 were designed (Table 8). 5ng of purified cDNA were tagmented in the presence of 5× TTBL and TTE Mix V5, incubated at 55°C for 10 min, and terminated with 5× TS buffer.

**Table 8:**
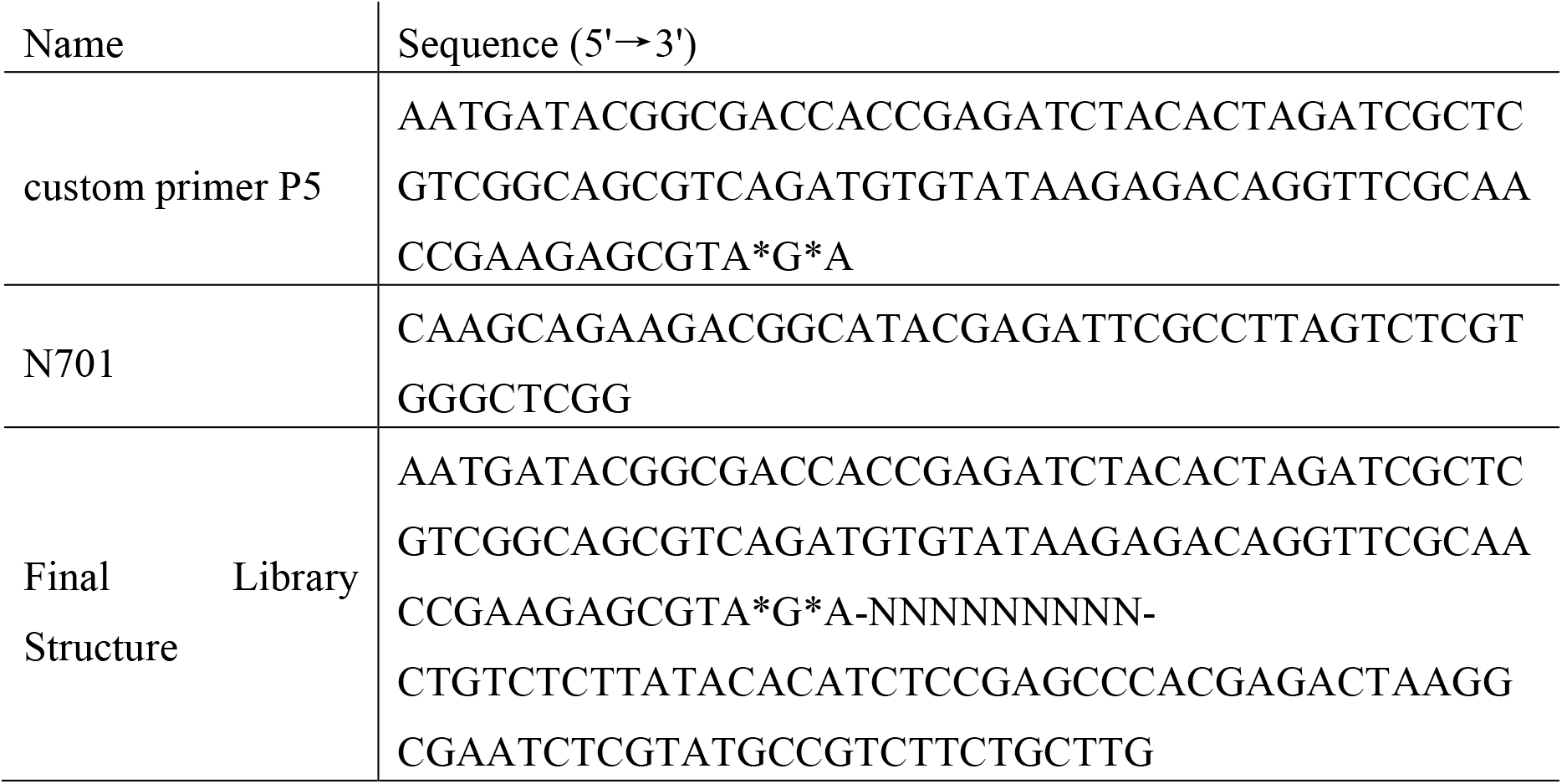
Sequence and Library Structure.

**Table 9:**
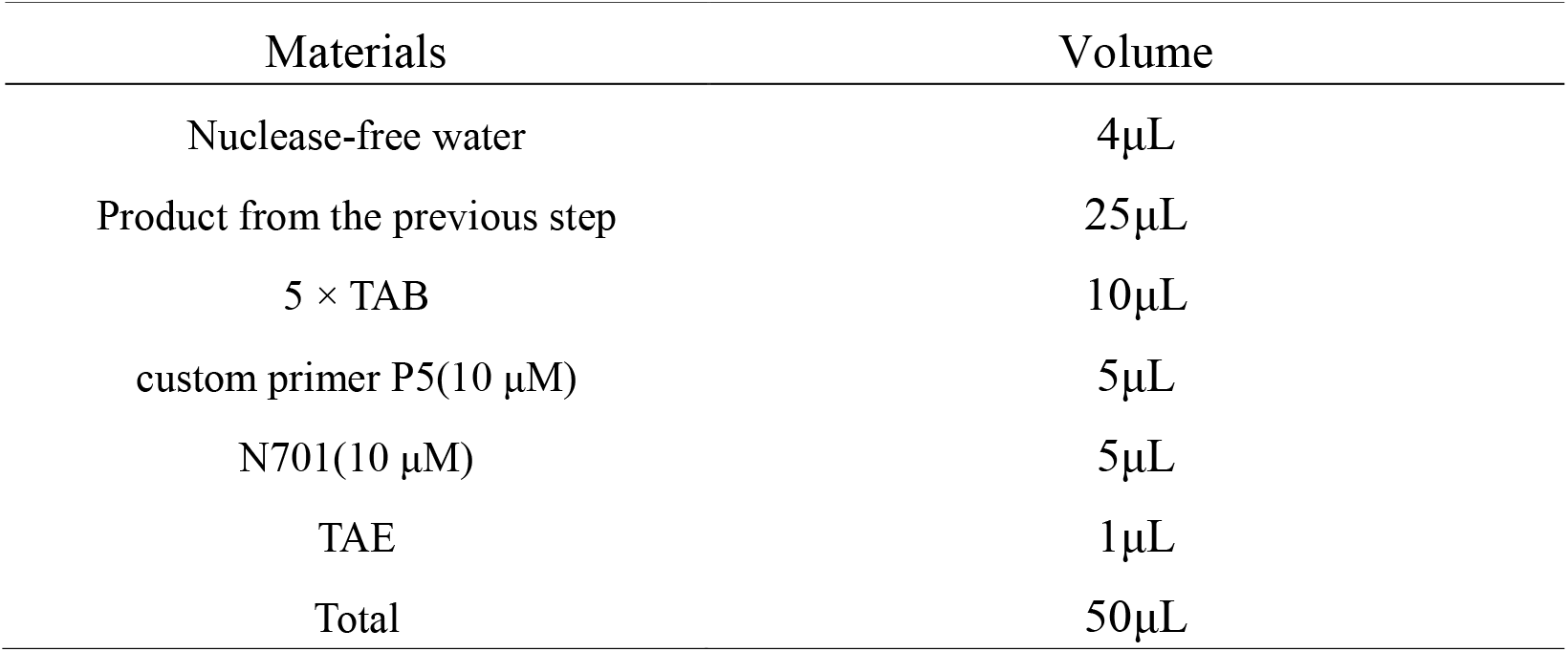
PCR Mixture.

| Materials | Volume |
| --- | --- |
| Nuclease-free water | 4μL |
| Product from the previous step | 25μL |
| 5 × TAB | 10μL |
| custom primer P5(10 μM) | 5μL |
| N701(10 μM) | 5μL |
| TAE | 1μL |
| Total | 50μL |

**Table 10:** PCR Program.

| cycles | Temperature | Time |
| --- | --- | --- |
| 1 cycle | 105°C | Heat lid |
| 1 cycles | 72°C | 3min |
| 1 cycles | 98°C | 30s |
| 12 cycles | 98°C | 15s |
|  | 60°C | 30s |
|  | 72°C | 30s |
| 1 cycle | 72°C | 5min |
| 1 cycle | 4°C | Hold |

PCR amplification of the library was performed using the mixture in Table 9, following the program in Table 10: an initial 72°C extension, denaturation, and 12 amplification cycles. Final products were purified using magnetic beads (0.7× ratio) and quantified before submission for Illumina sequencing.

## Results

### Microfluidic barcoding for single cell

The original concept of microfluidic barcoding existed for more than two decades when early microfluidic researchers attempted to create hybridization spots for nucleic acid detection with microfluidic channels. As shown in Figure 1, two sets of orthogonally arranged microfluidic channels (x and y directions) are able to general a two-dimensional grid of feature spots on a flat substrate, e.g. glass. This very idea of orthogonally arranged input was indeed borrowed from the well-known random access memory (RAM) circuit design in microelectronics for memory chips, the gist of which is to create a logarithmic dependency between the number of controlled features (*F*), and the number of available inputs (*n*): *n* = 2 ×*logF*. The advantage of such design is the capability to use a linearly increasing number of inputs (fluid channel in microfluidics, or voltage signal for RAM) to generate an exponentially scaling features (hybridization spots for microfluidics, or memory units for RAM), leading to a fairly scalable device architecture. Such elegant design saw an exciting revival where Fan *et. al*. used this exact strategy to create deterministic barcoding for spatial transcriptomics. In our work, we posit that these deterministic microfluidic barcoding can be used to index the mRNA transcripts in single cells. We first spin-coated and patterned micro-well arrays (a diameter of 28 μm and a depth of 25 μm, which is just sufficient to accommodate a single cell) with negative photoresist SU-8 on a clean glass wafer. The total number of micro-wells in this study is 4900. Two separate PDMS slabs with paralleled microfluidic channels (70 for each) were aligned with the micro-wells on the glass and reversibly bonded, flushed with desired reagents, and peeled off consecutively. Each microchannel possessed a slight larger width (50 μm) than the diameter of the micro-well. DNA barcode molecules with sequence structure shown in Figure 2b were injected into the first microchannel set (barcode-A, x-direction) through vacuum suction from the outlet side. Probes will react with the newly functionalized glass surface on the bottom of the micro-well and remained linked permanently. For the second set of microchannels, another set of barcode molecules (barcode-B, y-direction), mixed with linker probes (detailed structures shown in Figure 2b) were injected and subsequently linked with the previous barcode-A sequences using T4-ligase. At this point, a deterministic, scalable, completely bead-free device with single-cell indexing and mRNA capturing probe array is completed.

In order to verify the immobilization and linkage efficiency of the molecular barcodes, barcode-A were synthesized with FAM fluorescence (green) tags, whereas Cy5 fluorescence (red) tags were modified on barcode-B molecules. As shown in Figure 3a, after the immobilization of molecular barcode A was completed and the micro-wells were treated with sodium borohydride solution for reduction, the micro-wells were placed under a microscope and observed with green excitation light (488 nm). Compared to the other micro-wells, the micro-wells that had been introduced with barcode-A exhibited distinct fluorescence, indicating successful immobilization at the bottom of the micro-well. As shown in Figure 3b, after the barcode B linkage experiment was completed and the micro-wells were treated with SSC solution, the micro-wells were observed with red excitation light (552 nm). Fluorescence signal from barcode-B was only visible in the micro-wells that had been pre-loaded with barcode A, confirming the successful linking of molecular barcode B and A. As shown in Figure 3c, the micro-wells treated with KOH were placed under a microscope and observed with red excitation light. Compared to Figure 3b, distinct fluorescence was still observed in the micro-wells after KOH treatment, further validating the feasibility of the molecular barcode immobilization and linking experiments.

**Figure 3:**
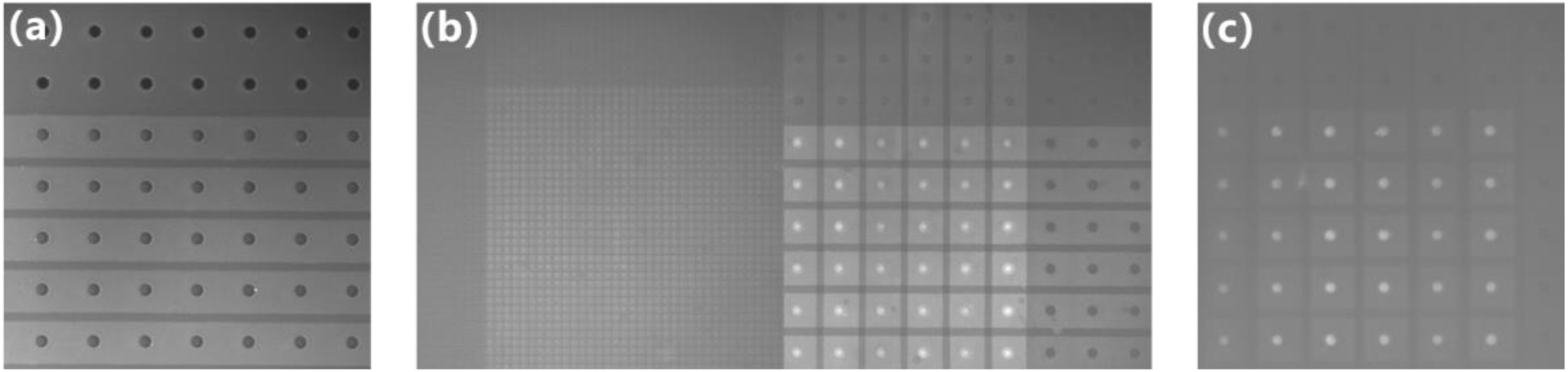
Results for molecular barcode immobilization and linkage experimental: (a) fluorescent image after barcode-A immobilization, (b) fluorescent image after barcode-B linkage, and (c) fluorescent image after KOH treatment.

### Single cell indexing

Next, we studied the effectiveness of the single cell indexing by loading the testing sample of K562 cells into the above platform. The beauty of this approach is that the trapping and indexing of individual cells are no longer even single Poisson distribution limited. Because we can carefully control the opening of the micro-well by photolithography such that only one cell can be trapped within. Therefore, after loading cell suspension, we can continue to maneuver the device until almost every cell fall into the micro-well, without much risk of duplets created.

A PDMS slab with microchannels was used for cell loading, featuring one pair of inlet and outlet interconnected by 10 paralleled channels, each of which can cover 7 rows of micro-wells on the glass substrate. This chip serves a dual function: first, it introduces the cell suspension onto the micro-wells region, allowing individual cells to sediment into the micro-wells under gravity. Second, it can then be used to introduce mineral oil mixed with lysis reagent, facilitating the oil-sealed lysis of single cells while preventing cross-contamination between micro-wells. In this experiment, cells were loaded into the chip through the inlet hole, and by adjusting the height of the tubing connection at the outlet, the pressure inside the chip could be conveniently controlled. This allowed the cells to be suspended right above the micro-wells, making it easier for the cells to fall into the micro-wells under gravity, as shown in Figure 4a. The cells were observed falling into the micro-wells under bright-field microscopy, as shown in Figure 4b. We also pre-stained the cells with Calcein dye before being loaded, making it easy to observe using the fluorescence, as shown in Figure 4c. The cell capture rate were calculated to assess the experimental performance:

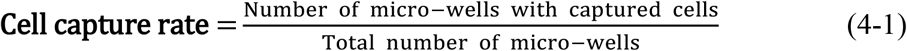

**Figure 4.**
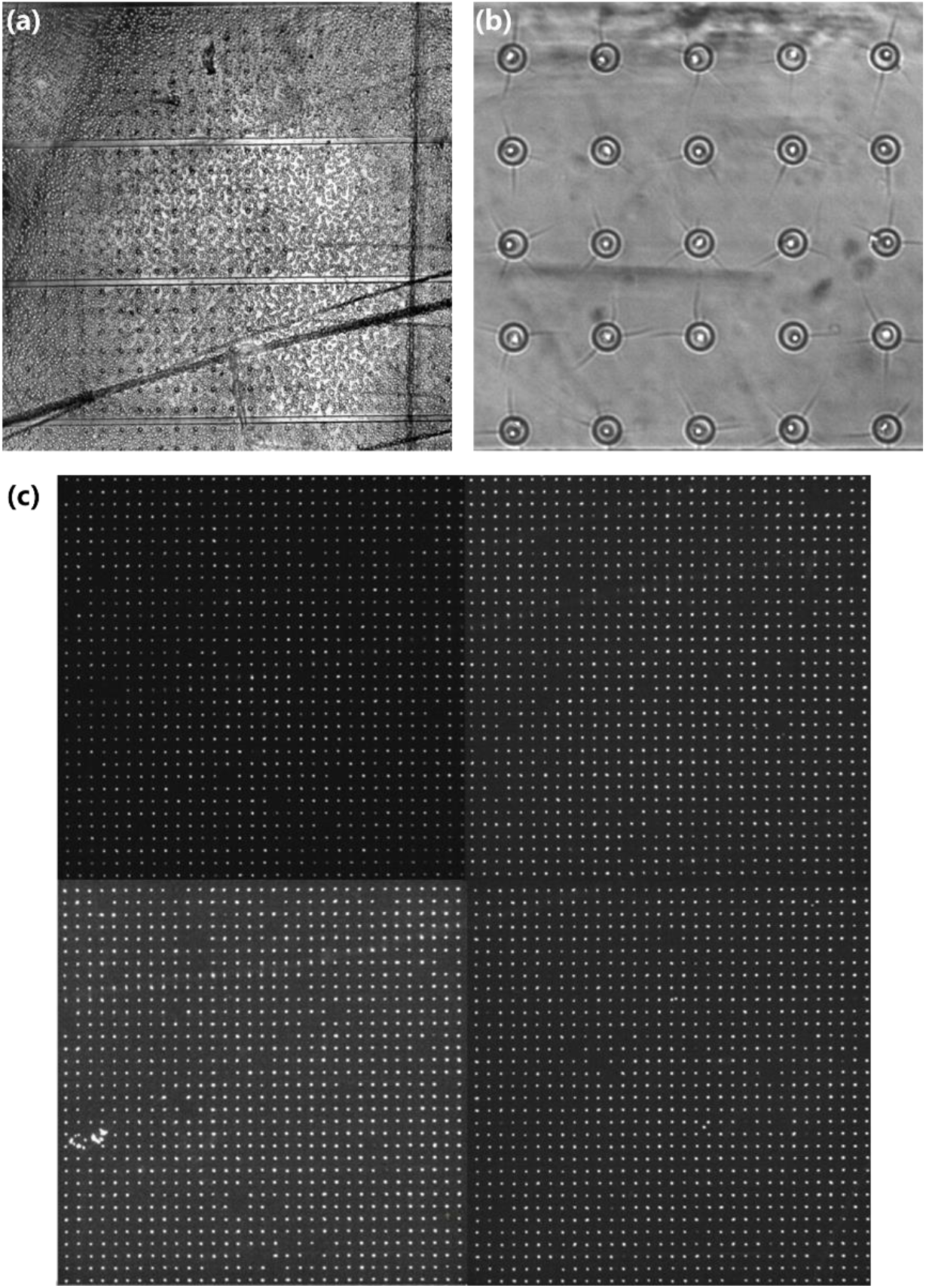
Micrograph for single cell capture under (a, b) bright-field and (c) fluorescence microscopy.

In this work, our device design was set 4900 (=70×70) total micro-wells on the platform, and the single-cell capture rate in this method can reach 96.6±0.2%. As microfluidic barcoding is inherently a deterministic process, the successful single cell indexing rate depends solely on the trapping rate of our platform.

### Single-cell mRNA sequencing

To demonstrate the utility of our platform, we first tested single-cell mRNA sequencing at smaller scale with 100 K562 cells. This requires 10 unique barcoding sequences each for the x-and y-direction. Figure 4a-b shows the proper trapping of 100 individual cells in the micro-well arrays that has gone through successful microfluidic barcoding. The device was then flushed with oil that contained surfactant nanoparticles. Upon fully filling the device with individual cells sealed underneath, lysis-agent would gradual precipitate and release itself into the aqueous solution inside the micro-wells, starting the actual lysis process.

By observing whether the micro-well is filled with fluorescence, it can be determined if the cells have been successfully lysed. As shown in Figure 5a, the comparison between bright-field and fluorescence-field images during the oil seal process shows that there is no difference in the fluorescence of cells before and after the oil seal, indicating that the oil sealing process does not affect cell viability or cause cell lysis or death. In Figure 5b, the comparison between bright-field and fluorescence-field images during the cell lysis process shows that unlysed cells (red arrows) retain their complete cell morphology in the bright field, and the fluorescence is concentrated within the cell, with the fluorescence region matching the size of the cell. On the other hand, lysed cells (green arrows) no longer show a complete cell morphology in the bright field, and the fluorescent signal spread the entire micro-well and was confined within the micro-well, validating the effectiveness of the oil seal and lysis methods used in this study. In Figure 5c, the comparison between the fluorescence images before and after cell lysis shows that after lysis, the fluorescence is evenly distributed throughout the micro-well and we indeed observed >99% of the cells have been lysed, further confirming the effectiveness of the lysis method used in this study.

**Figure 5.**
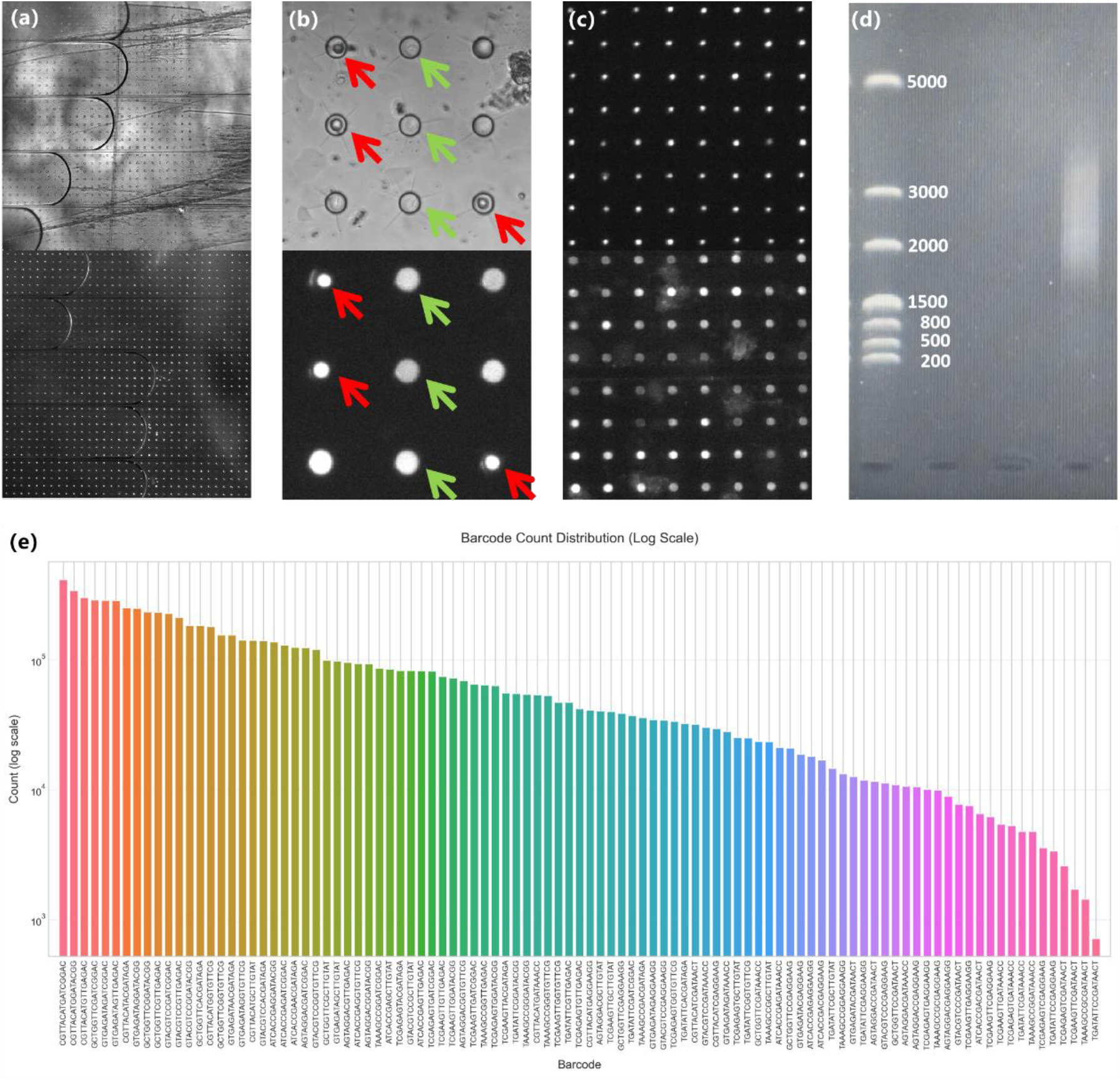
Cell Lysis Experiment Results (a) Comparison of Bright-field and Fluorescence-field Images During the Oil Seal Process. (b) Comparison of Bright-field and Fluorescence-field Images During Cell Lysis Process. (c) Comparison of Fluorescence-field Images Before and After Complete Cell Lysis. (d) Gel electrophoresis results for the collected cDNA samples. (e) Hit distribution for the whole 100 possible barcodes.

After subsequent incubation, barcoded DNA probes would capture mRNA released from lysed cells, and the oil was then removed from the device. The hybrid DNA-RNA duplex then went through the standard molecular reactions including cDNA synthesis, template switching and further PCR runs. The obtained samples were subjected to agarose gel electrophoresis to observe the length distribution of the samples. The results are shown in Figure 5d. The sample lengths are concentrated above 1500 bp, which confirms that the molecular barcodes successfully captured the mRNA. This further supports the reliability of the entire experimental procedure. Finally, the resulted DNA samples were sent for NGS sequencing. To verify successful single cell indexing, we analyzed the sequencing data as follows: we first listed all 100 possible combination among the 20 x+y direction barcoding sequences (Table 1×2). We then tried to identify all transcripts that contained any one of them using alignment algorithm. Figure 5e shows the distribution of hits for each of the 100 possible combinatory barcodes, indicating the incorporation of all of them in the cell transcripts.

## Discussion and Outlook

We presented a single-cell transcriptomic platform based on microfluidic barcoding that is easy to operate, readily scalable, completely bead-free, and achieve as high as 96.6% single-cell indexing rate. We demonstrated a cell capturing and processing throughput of >4500 using our platform, making it on par with most mature microfluidic methods (in the range of 5,000-10,000). With the nature of exponential scalability of our barcoding method, it is fully possible to extend to beyond 10,000 processing throughput. More critically, our work here represent an important paradigm shift for microfluidic single-cell omics studies. Beads with molecular barcoding are the default choice for single-cell indexing for the majority of high throughput single-cell omics studies. The SCI strategy (or equivalently the split-pool method), or MARS-seq 2.0 (which further perform pooling of multiple cell-sorted well-plates), requires extensive well-plate usage and pipetting efforts to reach higher throughput. Our method still leverages the unique advantages of microfluidic in handling large number of cells in a massively paralleled manner, but completely get rid of the trouble of dealing with barcoding beads. As shown, the extra effort and cost for manufacturing such beads are totally avoided. More importantly, the issue with the notorious double Poisson distribution is completely muted, which leads to what was obtained in our demonstration of ultra-high single-cell indexing rate. We are continuing the effort to use this novel platform in analyzing biomedically relevant samples and the results will be shared in the near future.

